# A comparison of turn identification methods on high-frequency movement trajectories reveals potential comparability issues between studies of movement ecology

**DOI:** 10.1101/2025.04.13.647400

**Authors:** Stefan Popp, Edward A. Codling, Joseph D. Bailey, Anna Dornhaus

## Abstract

High-frequency animal tracks must often be subsampled to allow a simple analysis of the movement on the most meaningful scale for the respective study. One way of achieving this is to identify ‘biologically significant turns’, compared to heading changings caused by ‘noise’. Many ‘turn identification’ methods have been developed, but the accuracy and consistency of such methods have rarely been validated against ground truth trajectories with known ’true’ turns and noise. We analyze simulated tracks with known parameters as well as two empirical tracks and identify turns with 10 different frequently used resampling methods. We assess the specificity and sensitivity of identifying the location of turns and compare the known mean step length and turn angle of the paths with the resampled trajectories. We found great accuracy differences between, and sometimes within, methods, even on simulated tracks of the same characteristics. Results of some methods were also highly sensitive to the user-set threshold the method requires (e.g. max angle). Overall, the best-performing methods in this study were DP and MRPA, methods used in human mobility research, and TPA, which is mostly used in primate research. We thus advise caution when comparing results of studies using different resampling methods and recommend justifying the use of the resampling method in addition to quantifying the sensitivity of results to the threshold value. This study is also an appeal to authors of novel turn identification methods to consider thorough comparisons in different scenarios with a wide range of previous methods, including those developed outside the movement ecology discipline.

## Background

The field of movement ecology aims to describe the movement of animals and link it to, among others, environmental, behavioral, and physiological characteristics and processes [1], helping us understand the biology of the investigated animals. Typically, continuous animal movements are recorded at a constant frequency, such as through video recordings (e.g. 25 frames per second) or tagging (e.g. GPS or ARGOS tracking with 1 location fix every hour or day) with recent technological advances allowing for the recording of accurate high frequency data (i.e. creating smooth movement tracks) over long periods of time [2–5]. While these highly detailed trajectories offer a wealth of data and allow for novel, more biologically insightful analyses than lower frequency trajectories [3], they also introduce new challenges outlined below: they amplify noise present in the movements and those introduced by the sampling method, and the sampling scale is finer than the scale of biologically meaningful movement.

For some applications, the most informative scale of movement is defined by ‘biologically significant’ turns, which usually correspond to a discrete decision of the animal to change its heading direction. Whilst some animals, such as dung beetles, are known to make reorientations at specific times through their movement trajectory [6], most are assumed to make continual adjustments to their movement in response to external stimuli, such as the perception of a food source [7,8], presence of a predator [9] or environmental conditions, such as path obstructions [10], noxious conditions [11] or landscape features [12]. In the absence of external stimuli, internal processes can also initiate turns, including untriggered stochastic neuronal firing [13], or patterns of gene expression [14]. Identifying biologically important turns is therefore an important question in ecology as it is known to have significant effects on important biological processes such as foraging strategies [15,16].

Noise across movement trajectories can be fine-scale navigational and locomotory variations in the movements, caused by the local environment conditions or individual mobility [17–19]. These small turn angles are uninformative for most research questions on the larger patterns and are not picked up in lower-resolution data. Noise can also be inaccuracies caused by the tracking method and technology [20–22]. The centroids of animals detected in video can move within the body outlines (‘optic wobble’), and telemetry data can have location errors of many kilometers [20]. Whilst various methods of dealing with noise caused by limitations of technologies have been developed [23–26], identifying key aspects of movement tracks (such as meaningful turns) through filtering out less relevant movement variations is a long-standing problem in movement analyses of both animals and humans.

Analyzing such high-throughput data with often used traditional step-turn methods, such as estimating step length and turn angle distributions, or path sinuosity, can lead to results that are not useful in terms of biological interpretation or comparison [5]. High-frequency sampling leads to turn angle- and acceleration distributions which are highly peaked about 0 because there is not much time for the animal to change the heading angle or speed between two fixes. The movement metrics at each point are thus highly correlated with their neighbors and contain little information about the behavioral patterns and decisions of the animal, as they become only apparent on a larger scale. To get meaningful and interpretable track descriptors, such as turn angle and step-length distributions, the points of high-frequency trajectories need to be subsampled or clustered in a way to match the scale of the movement process of the respective study. This could mean that if the animal makes a discrete decision on average every minute, the track should also have on average one point every minute.

One way of increasing the scale of a high-frequency track is subsampling, or resampling, where only certain points of the original trajectory are retained in the resampled track. This way, distances between points are often much larger than the noise, diminishing its effect on the analysis, whilst the retained turn angles reflect larger-scale patterns. Subsampling with a fixed interval is a simple technique which lowers the resolution of the data uniformly by retaining every *k*th point of the trajectory. However, as animals are not expected to perform true turns with a predictable period, sampling at equal time intervals could miss some important characteristics of the movement. If the sampling frequency is too low, the animal performs turns between the sampling points which are thus missed, and which leads to potentially missed turns and turn biases, wider turn angle distributions, and too short tracks [27–30]. Too high sampling frequency still retains many points which add little information about the biology of the animal but may instead keep small, biologically insignificant ‘noisy’ location changes. If the resulting interval of positional fixes does not match the approximate time scale of the behavior, there will be mischaracterization of movement, such as misidentification of travel and search states, and reduced inter-study comparability [31–34].

To identify biologically meaningful turning points, several turn identification (or ‘turn selection’) methods have been developed which subsample the track at irregular intervals based on decision rules about some geometric characteristics of the track. These methods are separate from behavioral state change models like HMM or BCPA, which infer behavioral states (like ‘search’ and ‘travel’) from the movement characteristics [5]. While turn identification methods tend to be designed for a specific type of movement or sampling scenario, it is not clear whether they also perform better for the scenarios they were conceived for than other methods. All turn identification methods require a threshold parameter(s) to be chosen a priori which determines the sensitivity of the algorithm, i.e. how many points are retained in the resampled track. This should be chosen so as to retain as much informative information about the movement track whilst removing extraneous information [5]. In movement studies, deciding the precise threshold value to use is often justified by reporting that changing the parameter(s) within some value range does not change the conclusions of the paper.

While such sampling methods have been routinely used for a long time [35], there is considerable debate about which method is best for finding true turns [31] and it is not fully understood when and if applying different methods would lead to different conclusions of a study. Despite this, there is a lack of direct comparison between competing methods. This is different in the fields of human mobility and computer science, where >100 resampling methods (here called ‘curve simplification’, ‘line simplification’, or ‘trajectory compression’) have been developed [36], compared to the roughly 10 used in biology. In the field of human mobility and traffic research, the objective is often not to infer decisions of the moving agent, but to efficiently compress the raw trajectories retaining the least points necessary that maximize the retention of important trajectory information (such as its geometry). Therefore, resampled tracks are here typically compared with the ‘noisy’ raw trajectories, regardless of where ‘true turns’ are thought to have occurred. Despite the fields’ similarity, there is a stark segregation between (animal) movement ecology and human mobility, with sampling or trajectory compression being a demonstrable area of potential, but unrealized, crossover. This potential information transfer therefore implies the need for closer integration between the fields of human movement and animal movement as discussed in Miller *et al*., 2019 [2].

Here, we aim to improve the use of sampling methods in movement ecology by highlighting the differences in outcome based on the choice of method. We specifically compare the sensitivity and specificity of a range of resampling methods to identify turns in raw tracking data of simulated agents which move in discrete steps and turns with varying levels of navigational noise. We seek to demonstrate that the choice of resampling method matters for robust and replicable studies and give indications for which method is best used in certain scenarios.

## Methods

### Overview

Our process is outlined in Figure 1, featuring four steps. A) To simulate high-frequency telemetry or video tracking data with known properties, we first generated artificial discrete-time ‘ground truth’ tracks. These tracks represented noisy straight-line movement interspersed with periodic reorientations (‘true turns’), mimicking a correlated random walk (see ‘Simulations’ for details). Variation in heading or turn angles at each point between true turns modelled biologically insignificant movements (‘noise’). B) These ‘ground truth’ tracks were resampled using methods from Table 1, with a range of threshold parameter values (see Table S3). We analyzed the resampled paths to measure c) how well each method identified true turns and avoided false detections using Receiver Operator Curves (ROC) and their areas under the curve (AUC). We also d) measured how well the resampled tracks retained the properties of the original tracks by comparing the mean step lengths and turn angles. This process was repeated for 14 scenarios of varying ground truth characteristics (see Table 2).

**Figure 1.**
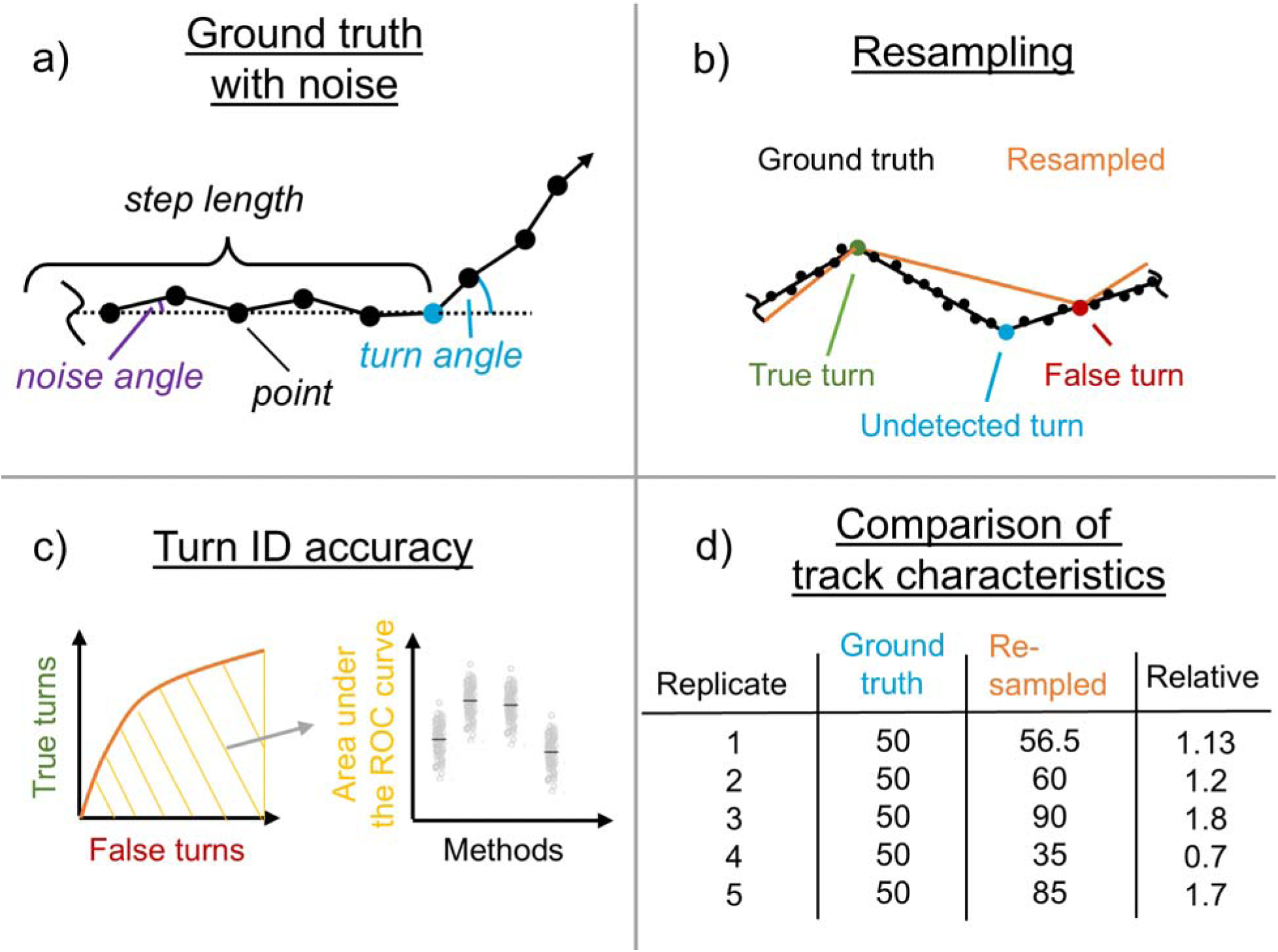
Methods overview. a) Ground truth track before noise. Blue dot is a ‘true’ turn. The step length is the distance between two true turns. Black dots are simulated sampling points 1 unit apart. Noise is introduced at each sampling point (not only at ‘true turns’). b) Resulting noisy ground truth tracks are resampled with the different methods, creating tracks which are composed of a new set of points. These ideally contain all ‘true’ turns and no falsely identified turns, which are far away from the next ground truth turn. c) Accuracy of resampling methods are assessed using the area under the Receiver Operating Characteristics (ROC) curve constructed by resampling the same ground truth tracks with different thresholds. Larger areas mean better true:false turn ratios across thresholds, i.e. a better tradeoff between sensitivity and specificity. d) The mean step lengths and turn angles of resampled tracks are compared with those of the ground truth tracks, validating whether the misidentification of turns leads to different track characteristics.

**Table 1.**
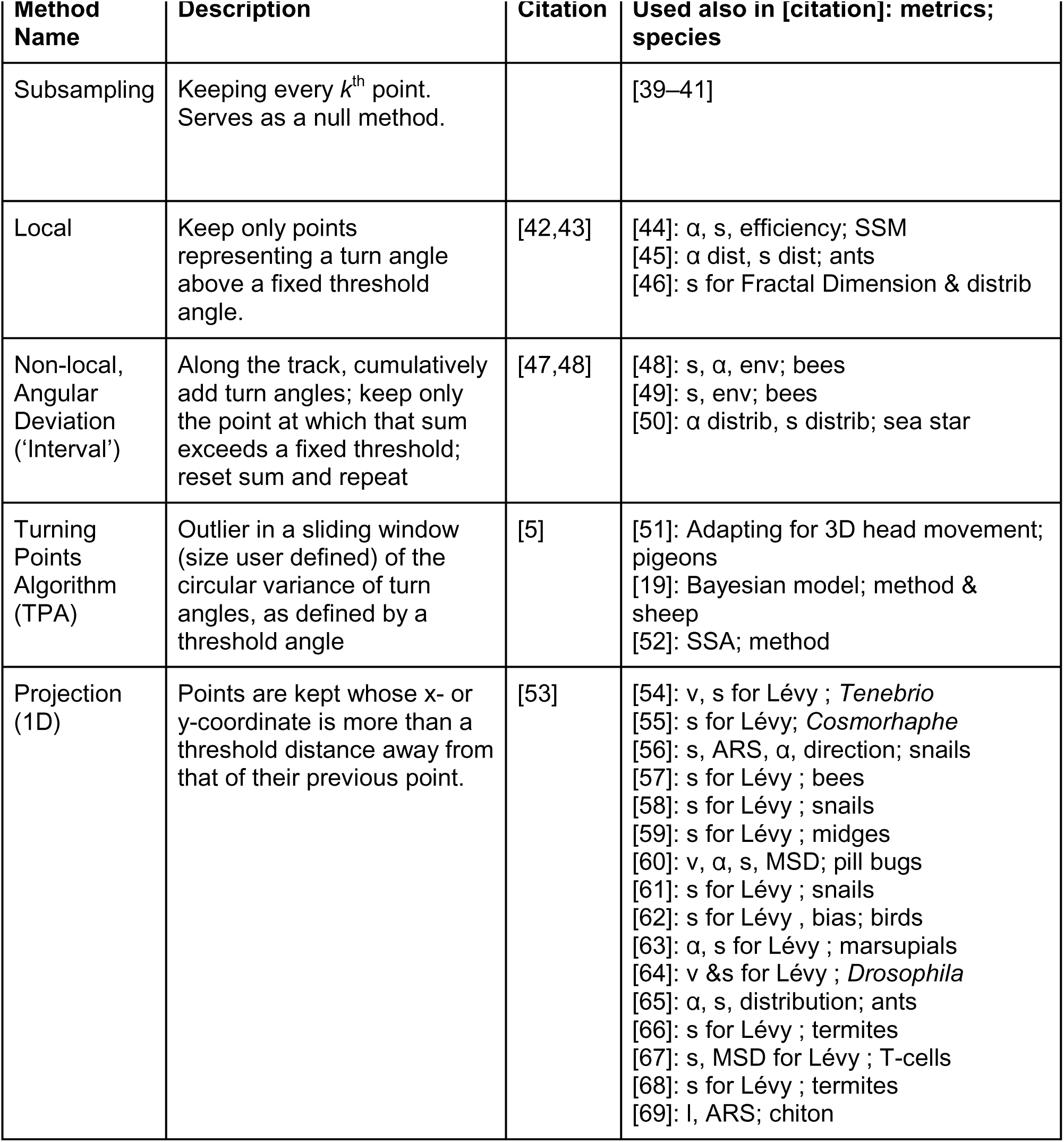
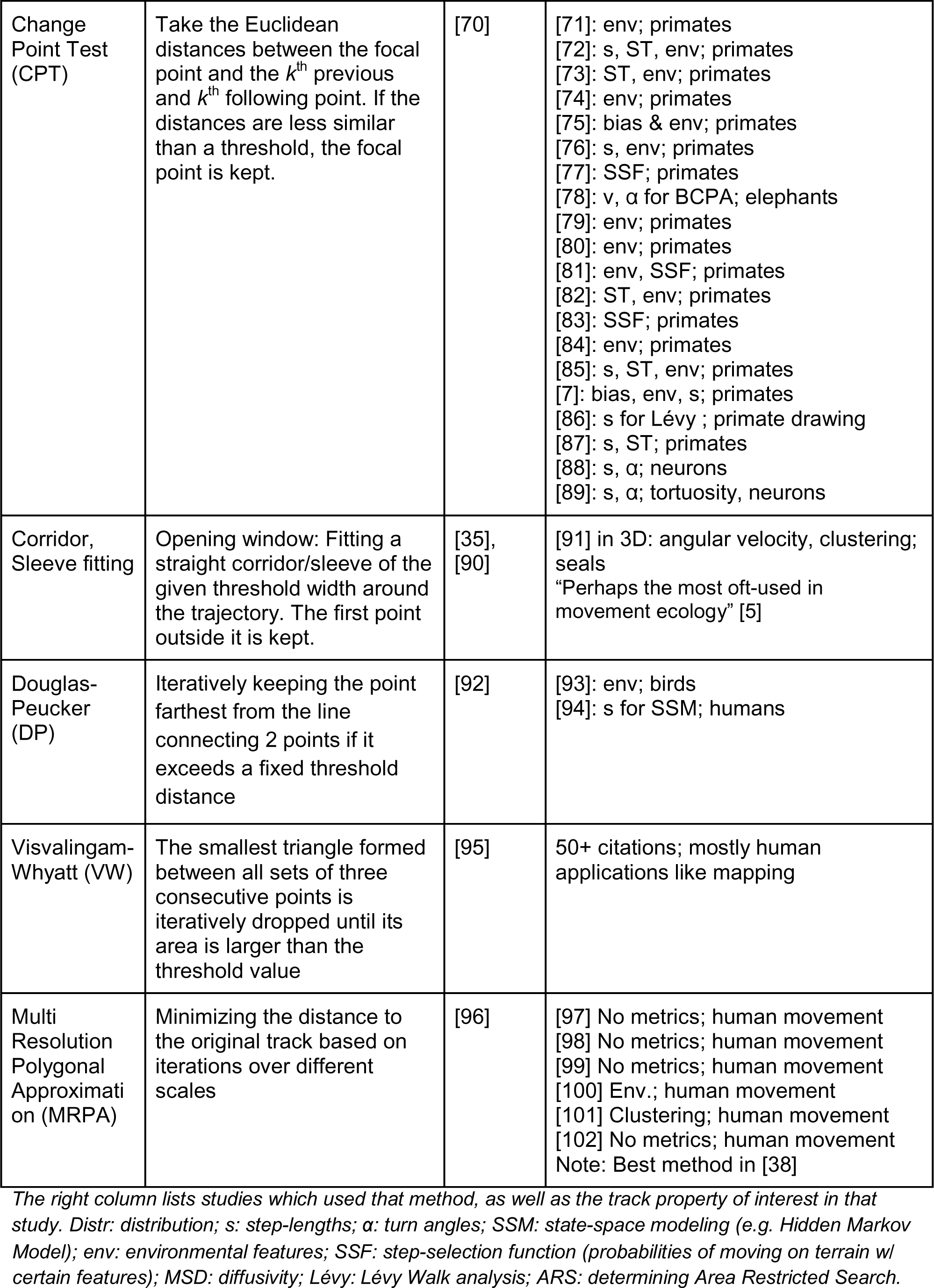
Resampling methods.

**Table 2:**
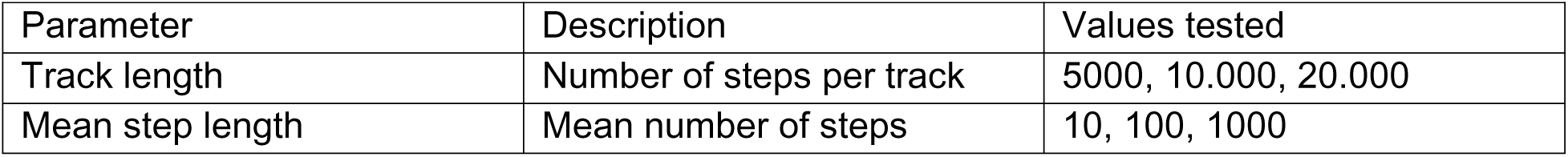
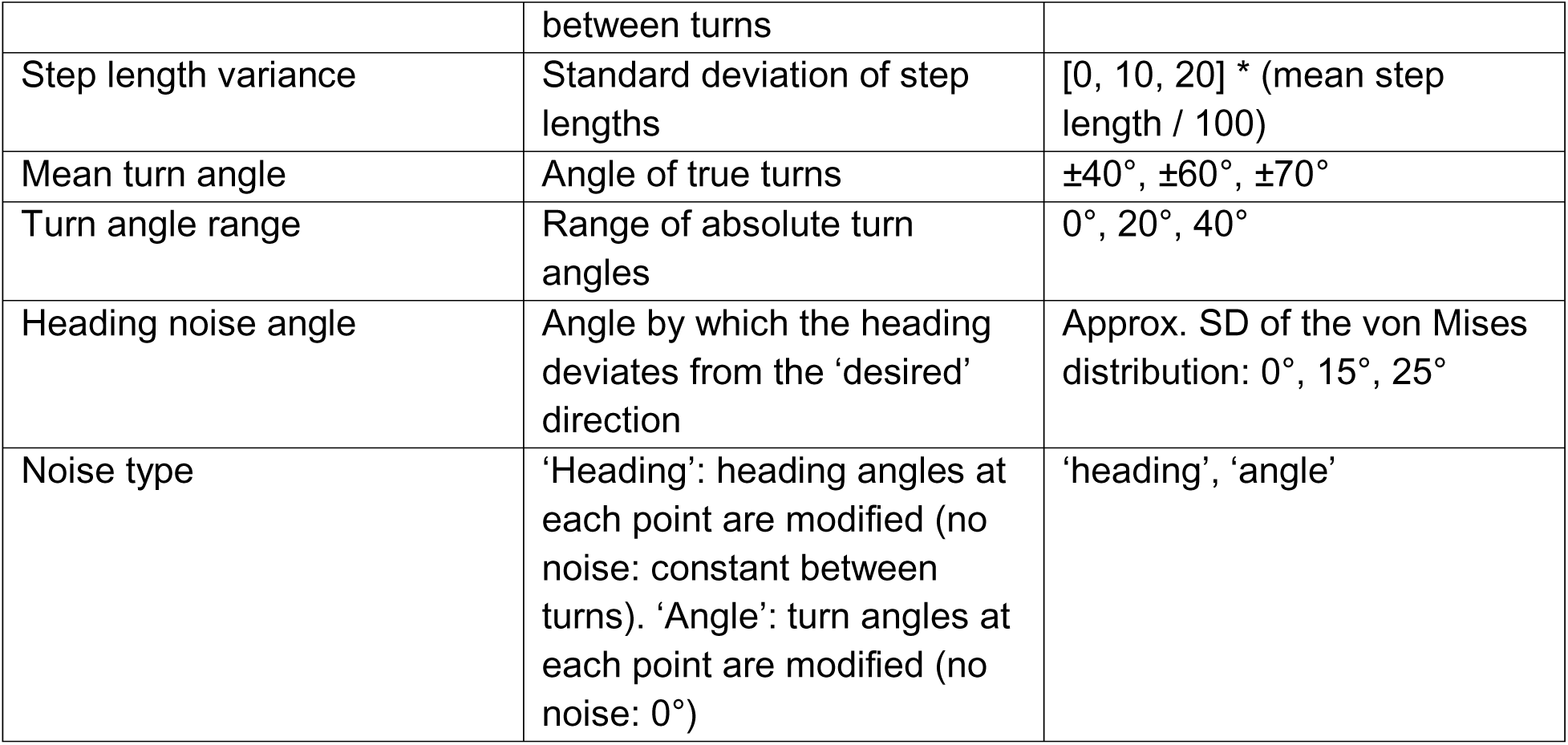
Parameters used to create tracks of different scenarios.

The resampling methods used are described in Table 1. These include resampling methods from the fields of animal movement, human mobility, and computer science. With methods chosen due to their popularity in the literature and, in the case of VW and MRPA, as they have been shown to have the smallest compression [36–38].

### Simulations

The purpose of this paper is to find out how well resampling methods can 1) identify where turns were made, and 2) not misidentify movement noise as being biologically significant turns, both together recreating tracks which resemble the original tracks without the noise. To assess how well resampling methods do this, we must thus know which turn angles in the track to be resampled are meaningful turns and which ones are noise. We simulate tracks for which we do have this knowledge and call them ‘ground truth’ tracks, allowing us to compare how closely the resampled tracks match the ground truth tracks without the noise. One could also hypothetically obtain ‘ground truth’ knowledge about movement tracks by ‘asking’ the movers where they made a turn decision, perhaps by reading certain brain signals correlated with this type of decision making. In this study, we simulated ground truth tracks (Figure S1) featuring l_P_ steps, with turns occurring every n steps, where n was drawn from a Gaussian distribution with its mean depending on the scenario, and standard deviation cr_n_ = 3. In the rare occasion of negative values of n, values were redrawn. Each turn angle, a_l_, was drawn from a uniform distribution with mean +µ_a_ , and range d_a_. The turn direction (i.e. sign of *µ*) for half the turns was randomly assigned to be left, and the other half right. We chose a uniform distribution over, e.g., a von-Mises-distribution, because we do not expect animals to concentrate their turn angles around one specific value, but rather perform turns that are appropriate for their current situation in the environment. At each point, a ‘noise’ heading angle (‘true’ turn or not) was drawn from a von Mises distribution with mean 0° and concentration parameter κ_c_. The concentration parameter is approximately equivalent to 1000_π ·_ (standard deviation of a Gaussian distribution)^-2^. To make this parameter more intuitive, we refer to the approximate equivalent standard deviation (SD) of the Gaussian distribution. For the ‘additive noise’ scenario, we instead added a noise *turn* angle using a von Mises distribution. The difference between these two noise types is that the noise in heading angle leads to straight lines between turns with local deviations from that path, while the noisy turn angle leads to slightly curved paths between turns but less local path jaggedness (Figure S1).

In our analysis, we focus mainly on 5 ‘scenarios’ with different characteristics which should correspond to different animal movements and sampling frequencies. Additionally, we perform sweeps across the parameters, the results of which we mostly report in the supplementary material.

All parameters were chosen such that the resulting tracks qualitatively resemble typical movement in animals across taxa, ecologies, and mode of movement [4,7,103,104] and are detailed in Table 2. 100 simulated tracks were generated for each of the 14 sets of parameter values.

### Resampling

All simulated tracks were resampled using the methods listed in Table 1. As each method requires a threshold or parameter to be chosen, the sampling of each ground truth track was carried out using 10 values (see Table S3), such that the ROC curves reached reasonably close to the 0,0 and 1,1 coordinates. The CPT method uses a second threshold which determines sensitivity and which we set to the recommended 0.05. The TPA method requires one angle- and one step-length (i.e. the sliding window size) based threshold to be set. In our main analysis we set the sliding window-based threshold to 2, which we beforehand determined to be the optimal value, but which is of course not known to the user in studies on animal tracks. We added additional analysis (S6) of the TPA method on the default scenario where the sliding window threshold is varied as well.

### Analysis

We conduct 2 types of analyses: the sensitivity and specificity of finding true turns in the scenario tracks, calculated with the Area Under the Receiver Operating Characteristic (AUROC) curve, and the difference of basic track characteristics between resampled and ground truth tracks in the parameter sweep tracks.

#### Turn identification accuracy

To obtain AUROC values, we followed the procedure laid out in [5]. First, we resampled the same ground truth track with a wide range of 10 threshold values specific to the methods. We then divided the track into segments of equal number of points (5 points for the default mean step length of 100 points, 3 and 50 points for mean step lengths of 10 and 1000 steps, respectively) and scored a true turn as being identified when the resampled track contains a point in the segment around the true turn. We calculated the proportion of ‘true turn’ segments which were correctly identified and the proportion of ‘false turns’, which are non-turn segments in which the resampling track contains a turn. With these two proportions, we constructed an ROC curve (Figures 2 & S5) where each point corresponds to one of the 10 thresholds used. Higher sensitivity and specificity correspond to higher y- and lower x-axis values, respectively. This means that the coordinate (0,1) in the ROC plot indicates perfect accuracy, and the distance between this point and a point on the ROC curve is a measure of performance of that specific threshold. In reality, the best threshold value is not known, which makes methods with relatively many threshold values near (0,1) more attractive. We then calculated the area under this curve from 0 to 1, which is largely determined by the best possible performance of that method. If methods did not reach either of these extreme values with any threshold, we connected the closest point to them with a straight line. We repeated this procedure for each of the 100 ground truth tracks per ground truth parameter set.

**Figure 2.**
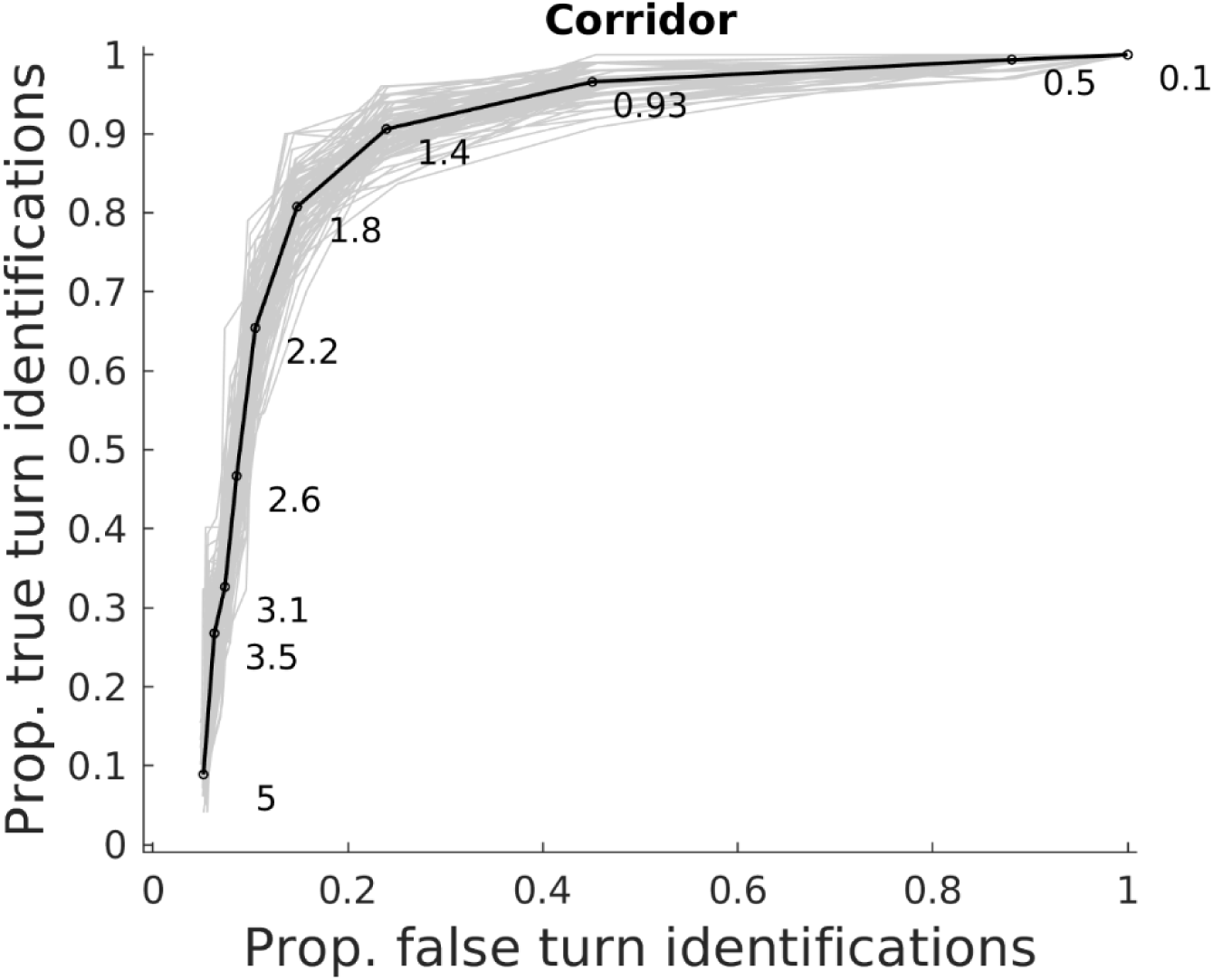
Example ROC curve of the Corridor method. Ground truth tracks were cut into segments of 5 points and segments around turns and non-turns counted which contained a resampled turn (true turns and false turns, respectively). The y-axis shows the proportion of ground truth turns identified and the x-axis shows how many non-turn segments contained a turn falsely indicated by the resampling method. The black line is the mean of the 100 ground truth tracks, shown in grey. Number labels are the threshold values for the Corridor method (the width of the ‘corridor’, outside of which a point is considered a turn point). The closer the curve comes to the top left corner, the better the method. The ground truth track scenario was ‘High noise’.

#### Comparison of track characteristics

Most studies on the movement of animals are not interested in finding the exact moment the animal made a biologically meaningful turn, but instead compare track metrics based on the step lengths and turn angles (see ‘metrics’ in Table 1). To test whether the differences in turn identification accuracy could change the characteristics of resampled tracks, the mean step length and mean turn angles of resampled tracks were additionally compared to ground truth tracks. These metrics were chosen as they are frequently used in movement ecology studies [39,105]. The relative accuracy of each track for each metric was then found by dividing the value of the metric from the sampled track by the value measured from the ground truth track (Fig 1d). We only included the resampled track of the best performing threshold for each method, as determined by the distance to the (0,1) coordinate in the ROC plot. We thus assume near optimal choice of the threshold by the user.

## Results

To simplify the comparison, we mainly show and discuss 5 exemplary scenarios: default parameters, small turn angle, small step length, large noise angle, and additive noise. Analysis of the other scenarios is provided in the Supplementary material. We chose these parameter variations to cover a wide (but non-exhaustive) range of animal movements and sampling regimes.

<u>Low accuracy of some methods in most scenarios, but small differences between scenarios</u> The methods Interval, 1D, Nonlocal, and VW generally had areas under the ROC curve of below 0.7, which was considerably less than the other tested methods (Figures 3, S5-S8). The consistently best performing methods were TPA, MRPA, and DP, consistently scoring a mean AUROC of above 0.9. DP was comparatively worse on ground truth tracks higher step lengths (i.e. more sampling points between turns), and MRPA and CPT were comparatively worse with shorter step lengths. While CPT had AUROCs of over 0.9 in the scenarios with heading angle noise added around straight-line steps, its accuracy dropped to around 0.8 in the scenario where noise turn angles were added (Figure 4, ‘Additive noise’), resulting in sometimes slightly curved steps between turns. Accuracies of all methods unsurprisingly declined with shorter step lengths, since this essentially means a lower sample size per point, and large noise angles (Figure 3, Shorter steps’ & ‘Large noise angle’). All other scenarios had minor effects on the accuracy of the tested resampling methods (see Figures S6-S8).

**Figure 3.**
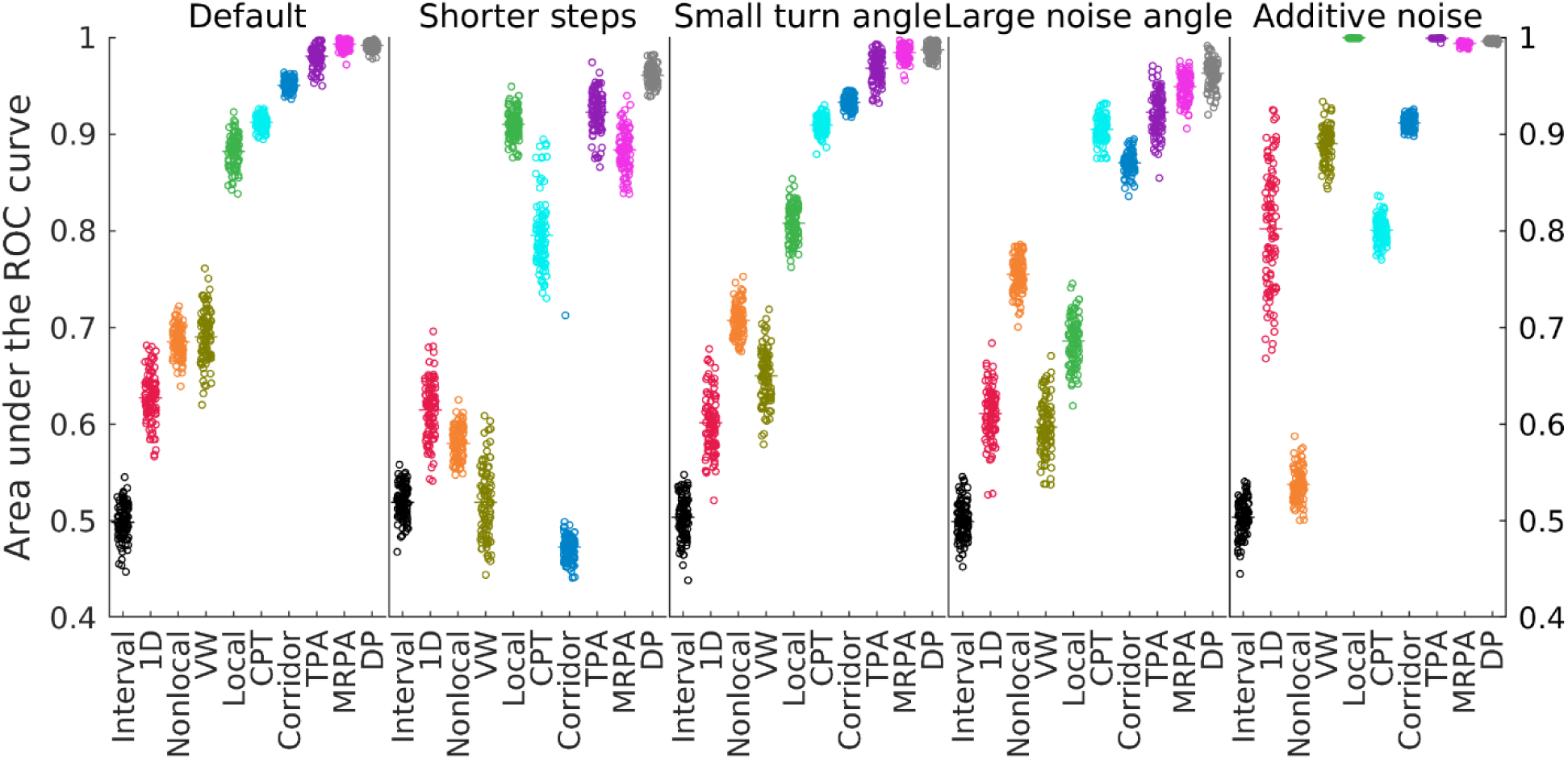
Areas under the ROC curves of example scenarios. Black horizontal lines are means and grey dots are resampled tracks of 100 replicate ground truth tracks per scenario. Larger numbers mean higher accuracy. The same data grouped by method are displayed in Figure S7.

**Figure 4.**
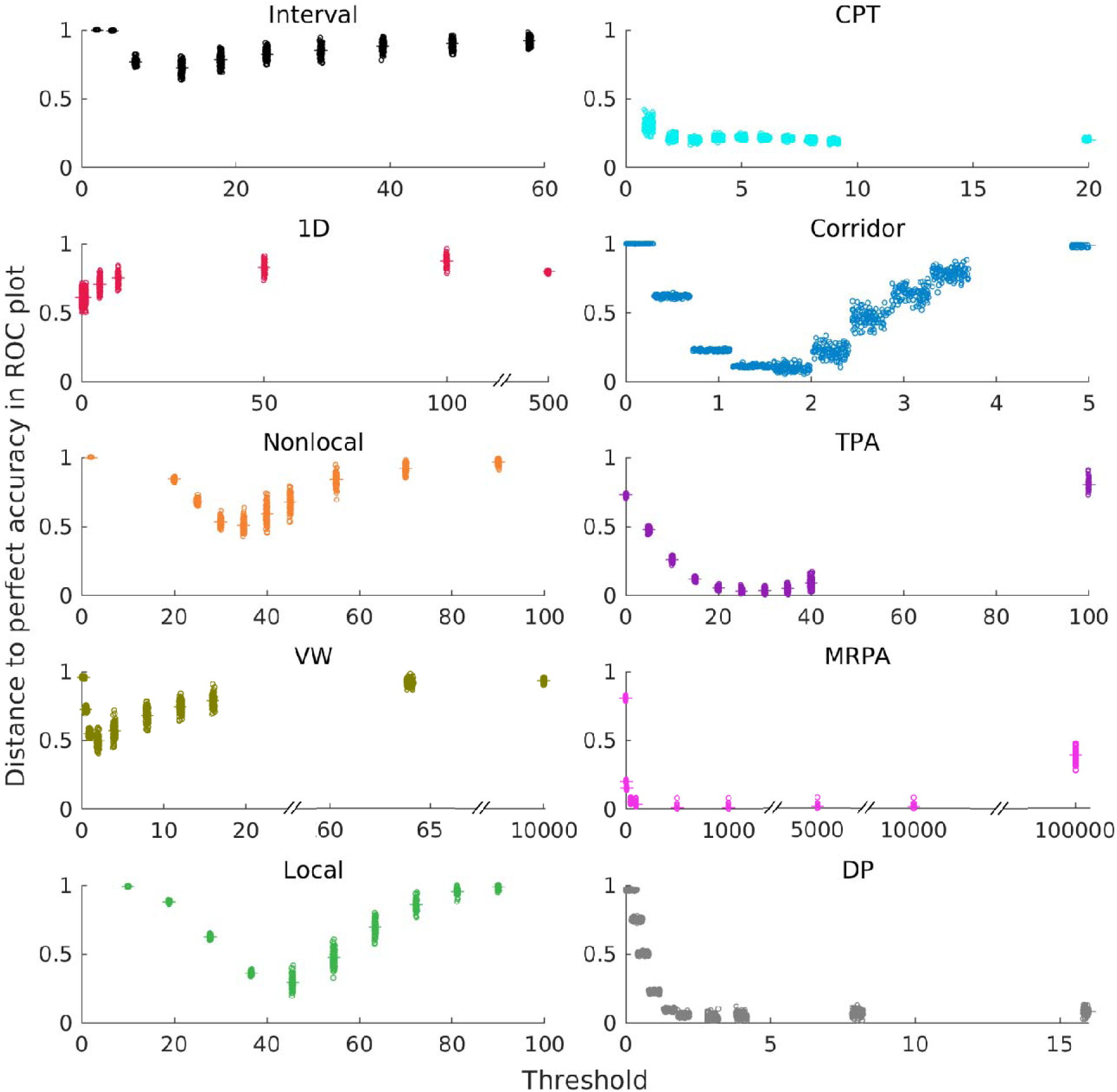
Accuracy depending on the threshold. The y-axis shows the Euclidean distance of the point in the ROC curve plot (Figure S5) to the upper left (0,1) corner, thus the distance to perfect accuracy of retaining only and all turns from the ground truth track (lower values are better). The x-axis shows the values of the thresholds required by the respective method. Black lines are means and grey dots are values for the 100 ground truth tracks of a scenario.

### Large variability with the threshold for some methods

For the Local, Nonlocal, and Corridor methods, accuracy can vary considerably when changing the thresholds slightly (Figure 4 and see the data labels in Figure 2 and S5 for threshold values on the ROC curves). To compare between thresholds, we define accuracy here as the Euclidean distance between the point resulting from a given threshold in the ROC curve to the top-left corner, i.e. perfect performance of finding all turns without false positives. For the VW method, this distance increases from 0.5 to 0.72 when changing the threshold from 2 to 0.5. The 1D and CPT methods draw short ROC curves, and thus cover a small space in the sensitivity-specificity tradeoff. The three best performing methods, TPA, DP, and MRPA were consistently biased toward being sensitive (i.e. identifying all true turns), meaning that turns were only missed with thresholds orders of magnitude larger than the other thresholds for whom the level of specificity (i.e. how few false identifications are made) varies with the threshold. The results shown here for the TPA method were obtained with the window size threshold set to 2, which is the optimal value determined in preliminary analysis. When varying the window size, the area under the curve value for the default scenario drops by up to 12% to 0.86 with a window size of 8, which is worse than the other 5 well-performing methods.

### Poor conservation of track characteristics with most resampling methods and large variation for lower step length scenario

To test whether the inaccuracies in finding turn angles result in dramatically different track characteristics, we compared the track characteristics ‘mean step length’ and ‘mean turn angle’ between the ground truth and resampled tracks. These metrics are often used in the literature to describe and compare tracks (see Table 1). We thus calculated how close these two metrics of the resampled tracks are to those of the ground truth tracks (accuracy in Figure 5 and variance in Figure S10). For both metrics, only MRPA come close to retaining the ground truth track characteristics across scenarios. For most other methods, the mean step lengths and mean turn angles of resampled tracks are 20% to over 30% larger or smaller than those of the ground truth tracks. Similar results as for the mean step length were found for the mean absolute turn angle. (Figures 5). Additionally, there is great variation in the accuracy in the scenario with shorter step lengths (i.e., fewer sampling points between true turns; Figure S10).

**Figure 5.**
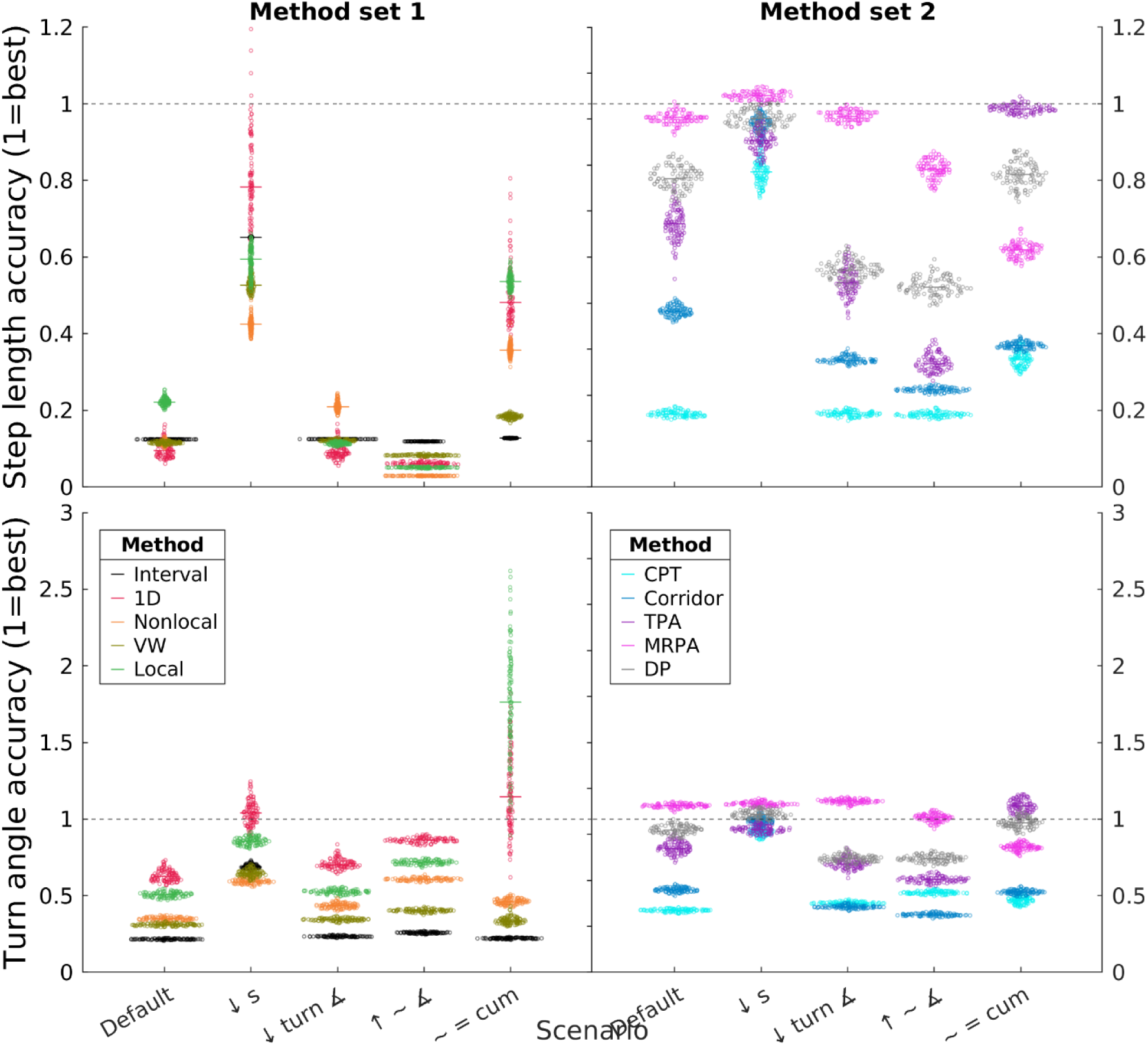
Resampled track mean step length and turn angle accuracy. The x-axis shows the scenarios shown in the previous figures. Different colors correspond to the different resampling methods. Left and right panels each show one half of the resampling methods for clarity; note different axis scales. All data are from resampled tracks resulting from the thresholds yielding the best turn identification accuracy as defined by the point which is closest to the (0,1) corner of the ROC curve, for the respective method. Top row: Accuracy of mean step length estimations. Bottom row: Accuracy of mean turn angle estimations. A value of 1.1 means resampled tracks overestimate the mean step length by 10%, and a value of 0.9 indicates underestimation by 10%. Horizontal dashed lines at 1 indicate the value of perfect accuracy. Points are outcomes for individual tracks (x-axis jitter added; all points in a ‘column’ represent resampling outcomes from ground truths with the same mean turn angle), and lines are medians.

### Empirical data

To show whether the results of the inconsistent track metrics are qualitatively similar with real-world data, we repeated the resampling and calculation of the mean step length and mean turn angle on trajectories of different animals in freely available data sets (ant from [4], cat from [106]; Figure 6, right column). The cat track might most resemble our ‘Small turn angle’ scenario, while the ant track is probably close to a combination of ‘Large turn angle’ and ‘Short step length’. The ant data was gathered by filming an ant freely moving in an arena with 25 fps and tracks were subsequently subsampled to contain points with 2 mm distance to the preceding point, eliminating most noise due to the tracking method. The positions of the cat were taken with GPS with location error of ca. 3.3 m every 20 min between 17:00 h and 9:00 h and every 96 min in the other time, and in subsequent data cleaning outlier fixes were removed. We only compared step and turn metrics between resampled tracks, since the ground-truth movement process, and thus existence and location of ‘true turns’ are unknown. We cut both raw trajectories to 1000 points for comparability.

**Figure 6.**
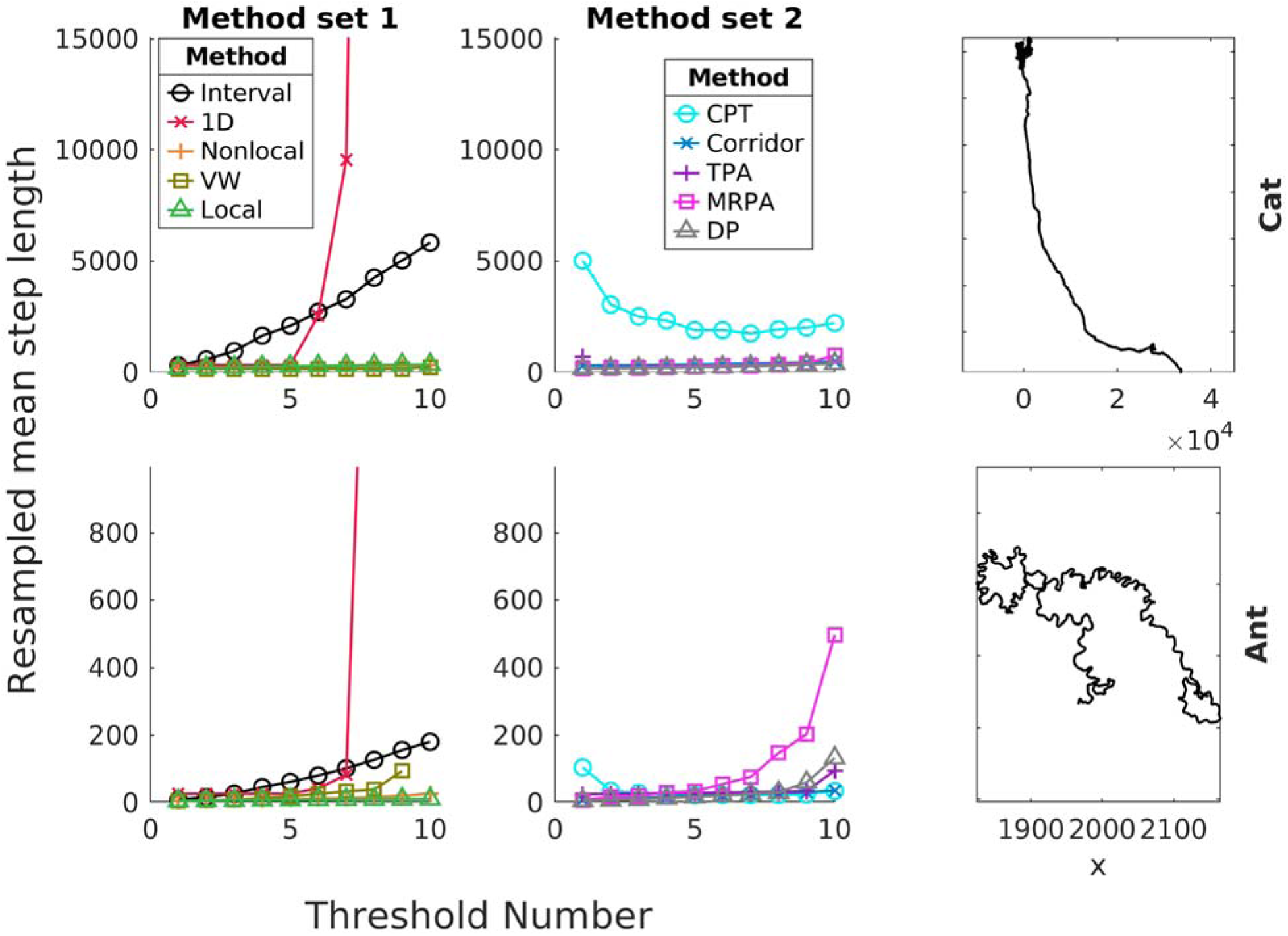
Empirical tracks and results. Top row: a feral domesticated cat in Australia, from (from [106]), bottom row: a Temnothorax rugatulus ant from [4]. Left and middle panels show mean step lengths of resampled tracks. The y-axis shows the mean turn angle of resampled tracks and the x-axis shows the ordinal number of the method parameter used (see table S3 for real threshold values). Right panels show the empirical tracks used.

## Results – Empirical

### Variation between methods when resampling empirical tracks

There was also considerable variation in metric values between methods and threshold values when resampling real (empirically measured animal movement) tracks (Figure 6). The mean step length of the resampled tracks varied by over 10% between some methods for medium threshold values, although there often exist threshold value combinations which result in similar track characteristics.

## Discussion

### Summary of results

We showed that methods of resampling a movement track to retain only the ‘biologically significant’ turns are often not very accurate and that there is great variation between these methods. As expected, resampling accuracy also depends on the optimal choice of threshold, which is unknown in real-world scenarios. Some methods additionally show large accuracy variability when applied to different tracks of the same characteristics. The inaccuracies and variation between and within methods result in tracks of highly inconsistent turn angle and step length characteristics. With few exceptions, methods perform generally consistent relative to each other independent of ground truth (i.e. animal) track characteristics.

### Method accuracies for finding turning points

We defined the accuracy of the resampling methods with the area under the ROC curve, which was built by plotting the proportion of identified true turns *versus* false positive turns for a wide range of method thresholds. Across the scenarios we consider here, a somewhat clear ranking of resampling method accuracy emerged. The Interval, Nonlocal, 1D, and VW methods all performed relatively poorly. Local, CPT, and Corridor typically perform well, with AUROC values of over 0.9. The overall best performing methods in our study were TPA, DP and MRPA, with AUROC values of over 0.98. A notable exception is the top performance of Local in the ‘additive noise’ scenario, where ground truth tracks contained turn angle noise between turns instead of heading angle noise, leading to smooth curves between turns instead of jagged straight lines between turns.

### Method accuracies for retaining step and turn characteristics

Most movement ecological studies are interested in comparing the statistical movement characteristics of the tracks, instead of trying to identify the locations of the turn ‘decisions’. We show that also the two often used metrics of mean step length and mean turn angle vary widely between resampled tracks of different methods, and also within some methods. The patterns of track characteristics accuracy are similar to those of the ROC accuracies. MRPA, created tracks which usually matched the ground truth tracks well, i.e. with no more than 40% differences in values, with TPA scoring even better in the short steps and cumulative noise scenarios. The major differences to the ROC results are that CPT and Corridor perform among the poorest, with vastly underestimated mean step length across scenarios. This inconsistency between methods may thus lead to different conclusions in studies using different methods. For example, if one study resampled a track which resembles our Cumulative Noise scenario using the Local method, it might analyze resampled tracks of 100° mean absolute turn angle, while another study resampling the same raw tracks using the Nonlocal method would base their further analyses on tracks of 25° mean absolute turn angle.

### Reason for differences between methods

We do not have an intuition for the ranking of methods we observe. Local, Nonlocal, and TPA use turn angles as basis for their algorithms, while 1D, CPT, DP, Corridor, and MRPA are based on Euclidean distances between the raw and resampled track, and VW uses areas of the track as threshold criterion. There does not seem to be a systematic correlation between the type of threshold used and the accuracy or variability of the method.

### Influence of ground truth characteristics

Surprisingly, changing most ground truth characteristics within the bounds explored here did not change the relative accuracy of resampling methods much. The number of points between turns (here termed ‘step length’) had among the greatest influence on resampling accuracy and variation, in terms of both identifying the location of turns and retaining the track characteristics. This shows the importance of gathering tracks, or track segments with high sampling frequency. This parameter, however, does not change the status of DP, MRPA, and TPA as being the most accurate resampling methods.

### Noise in the animal movement

We introduced two types of noise: deviations of heading angle and turn angle between turns. Heading angle noise (the default in our dataset) simulates local navigational or locomotory inaccuracies which are quickly corrected, while turn angle (‘cumulative’) noise leads to cumulative heading angle deviations and curvy steps between turns, which models animals which either drift off their original course or who do not have a fixed goal direction. When the heading noise is increased, both accuracy and variation within methods increase for most methods. When the noise between turns is turn angles, the TPA method performs best with near perfect accuracy, while the other methods maintain their accuracy levels.

### Thresholds

Translating the above results to the real-world application of resampling methods requires selecting the best threshold for the scenario. However, the optimal value of the threshold often requires already extensive knowledge about the movement mechanism and/or characteristics, leading to a biased selection of the threshold, leading to resampled tracks that ‘look correct’.

This issue has been discussed previously [31,107], although authors creating and using these methods either claim robustness to changing the threshold [43], or use some mathematical or biological reasoning to identify the optimal threshold [e.g. 53,55,115]. This dependence on selecting an appropriate threshold value is exacerbated when more than one threshold must be set, as in the TPA and CPT methods. For TPA, we set the second threshold (moving window size) to the previously determined optimal value, which in practice is not trivial to identify and which differed by an order of magnitude from those reported in the original TPA paper [5]. If this threshold was not optimally selected, accuracies dropped for some values far below those of the other two top performers. This method thus requires reliable knowledge about the relationship between the thresholds and track characteristics one intends to identify. In this study, the method most robust to the threshold selection was MRPA, which produced results within 10% optimal accuracy with a range of thresholds from 50 to far over 10,000. In this method, the decision power over which points to keep is shifted toward the algorithm itself, and less the threshold input. It is likely that this quality arises from the multi-scale nature of the method, which contrasts with the other, more locally operating methods considered here.

### Variation between ground truth tracks of the same scenario

Especially with ground truth tracks where few (10) locations between true turns were sampled, there is great variation in the tracks resampling different replicates of the same ground truth tracks. In studies on animals, this inconsistency potentially leads to spurious differences detected between mechanistically identical animal tracks. Thus, to achieve comparability between studies, tracks in different datasets or even species might have to be resampled with the same method and on the same (relative) scale [4]. There is furthermore the possibility that results of any individual study are not robust to which method is used, which is usually not selected based on a rigorous assessment of fit to the studied organism and context (see any ref in tab 1, ‘Also used in’). On the whole, MRPA performs better than the other methods, with DP and TPA being also very accurate. Local is a good method if the appropriate threshold angle can be guessed with high accuracy and there exists low turn angle variation.

### Empirical

Applying the resampling methods to empirical tracks of a cat and an ant results in potentially similar variation of mean step length and mean turn angle between methods as in the simulated tracks, depending on the chosen thresholds. With a certain combination of selected thresholds i.e., for some methods relatively high, and for others relatively low thresholds, one obtains similar track characteristics with many of the here tested methods. The characteristics of Interval, 1D, and CPT did not overlap with those of the other methods, regardless of the selected threshold. Future work may reanalyze data from published studies with different resampling methods to evaluate which conclusions change with a different use of method.

### Methods from other fields

Methods not used in biology, but in human mobility research should be considered in biology, since some of them perform better than some methods that are used in biology, notably MRPA, which achieved among the best performance and by far the lowest variance in our analysis [36,109,110]. Methods in these adjacent fields are often improved on and comprehensively compared among each other [36,38,107], which is facilitated using common benchmark datasets and the fact that the ideal result is known: retaining as much information while reducing the number of points as much as possible. In biology, one rarely knows how well one has achieved one’s aim, since the state of the animal will always have to be inferred from collected data, and discrete turn decisions may not even exist in the study species, as we simulated with the curvy tracks. Biologists should thus consider using promising methods developed outside of movement ecology or biology in general.

### Movement speed as an additional threshold

Another way of potentially improving the accuracy and replicability of resampling is to also take the speed (or acceleration) of the animal at each sampling point into account as an additional threshold. There are several methods which follow this approach (see S4 for overview) but were not included here due to the exponential increase of the number of simulation parameters, which would make a comprehensive analysis like we did here infeasible. Future work may compare the variability of results between these methods using a set of empirical data.

### Other ground truth track characteristics not considered

There may be other track characteristics or parameter values we did not include in this study which may influence the relative performance of the resampling methods. Empirical tracks could, for example, include noise due to obstacle avoidance, be composed of steps with exponential length distribution, or contain correlations between step and turn characteristics. They could also be created by different movement processes, such as composite random walks [111] or continuous turning [108].

### Curvy tracks

Most animal movement may be generated continuously, even though our sampling methods always sample the tracks in discrete steps and turns [112]. Since there is arguably no way of ‘correctly’ identifying individual points where entire turns occur, it might be more comparable and truer to the movement process of the animal to consider a different trajectory description framework, like the persistent turning walker model, which assumes smoothly changing angular velocities [5,113,114]. If necessary, such trajectories could thus be downsampled to segments of similar angular velocity, as defined by a threshold rule, thus retaining the curvy nature of the track [115].

## Conclusion

We show that the claim of identifying the location of biologically meaningful turns in animal tracks is not always true for some frequently used resampling methods in biology. These inaccuracies in turn identification resulted in different characteristics of resampled tracks, potentially leading to wrong conclusions of the studies that employ these methods. More likely, the differences in resampled tracks between methods and used thresholds leads to lower comparability of results between studies. However, since we found negligible differences in the relative performance of methods between ground truth tracks of different characteristics, we can give specific recommendations here. MRPA, which has to our knowledge never been used in ecological studies, consistently performed among the best, had low variability within sets of replicated ground truth tracks, and was very insensitive to the choice of the threshold. Although CPT and TPA also scored well in our study, the fact that 2 thresholds must be set speaks against the utilization by novel users. More generally, we advise justifying the use of the resampling method, and possibly testing the robustness of the results of the study to the choice of resampling method and its threshold. Authors of novel resampling methods should include an extensive test of their method on ground truth tracks with different (step) lengths, turn angles, noise types, and noise levels, as well as compare their output to that of the top performers in our study and possibly others used in the human mobility literature.

## List of Abbreviations

AUROC: Area under the Receiver Operating Curve
DP: Douglas-Peucker
CPT: Change Point Test
1D: 1-dimensional projection method
TPA: Turning Points Algorithm
VW: Visvalingam-Whyatt
MRPA: Multi-Resolution Polygonal Approximation

## Declarations

### Ethics approval and consent to participate

Not applicable

### Consent for publication

Not applicable

### Availability of data and materials

The datasets generated and analyzed during the current study are available in the OSF repository, https://osf.io/kjrnv/?view_only=6e2c7f4c6fd74f189f2c0e3c7e992c0a.

## Authors’ contributions

All: Conceptualization, Methodology; S.P. and A.D.: Project administration; S.P.: Software, Formal analysis, Investigation, Resources, Data curation, Writing – Original Draft, Visualization; A.D., E.C., and J.B.: Writing – Review and Editing; A.D.: Supervision, Funding acquisition

## Acknowledgements

We are grateful to Brian Enquist, Dan Papaj, and Tom Kennedy for helpful comments on the first draft of the manuscript.

