## Supplementary material for "A comparison of turn identification methods on high-frequency movement trajectories reveals potential comparability issues between studies of movement ecology": SI

### Methods

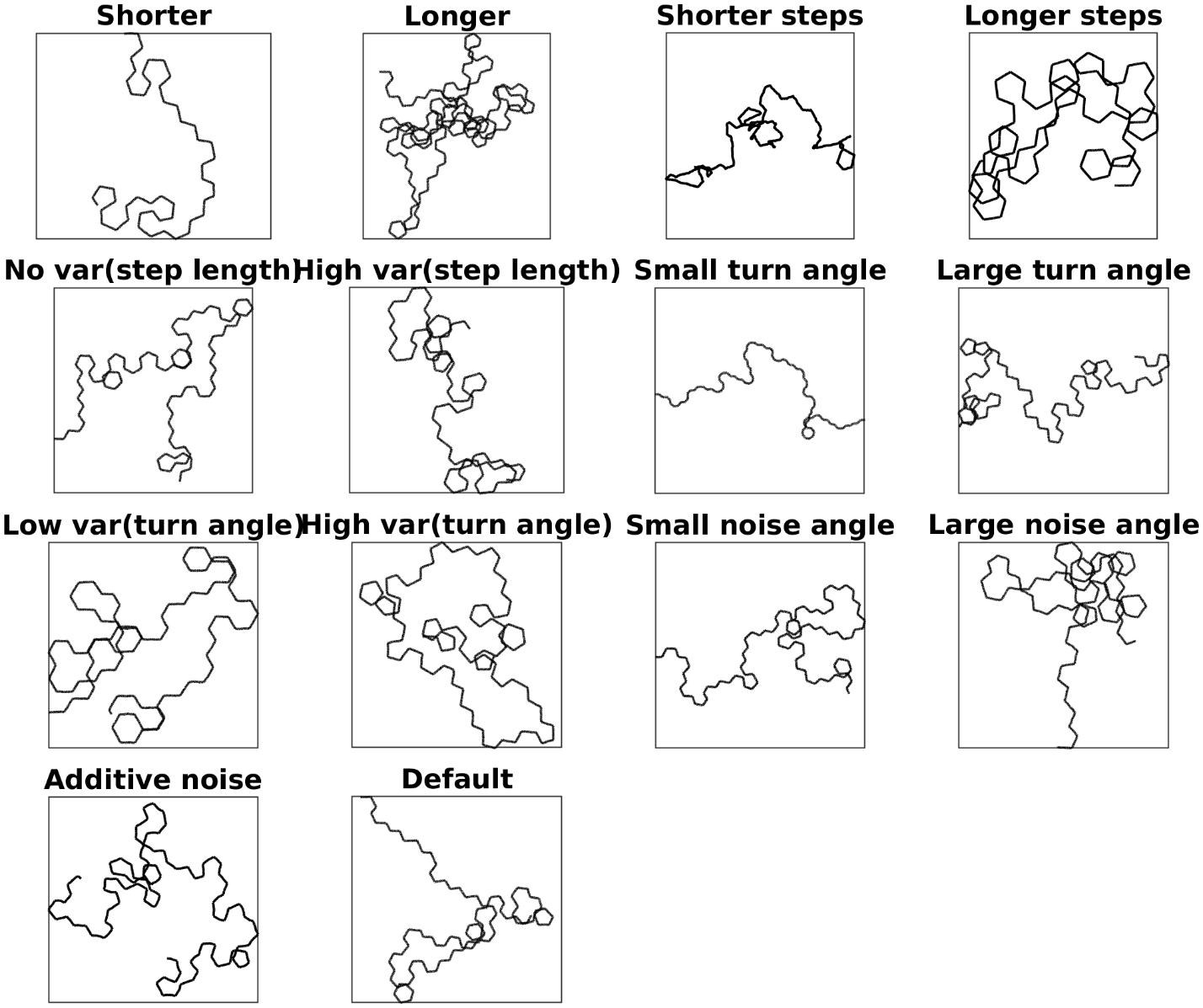

***Figure S1.*** ***Map representations of all simulated tracks.*** *Panel titles indicate how the tracks differ from the default track.*

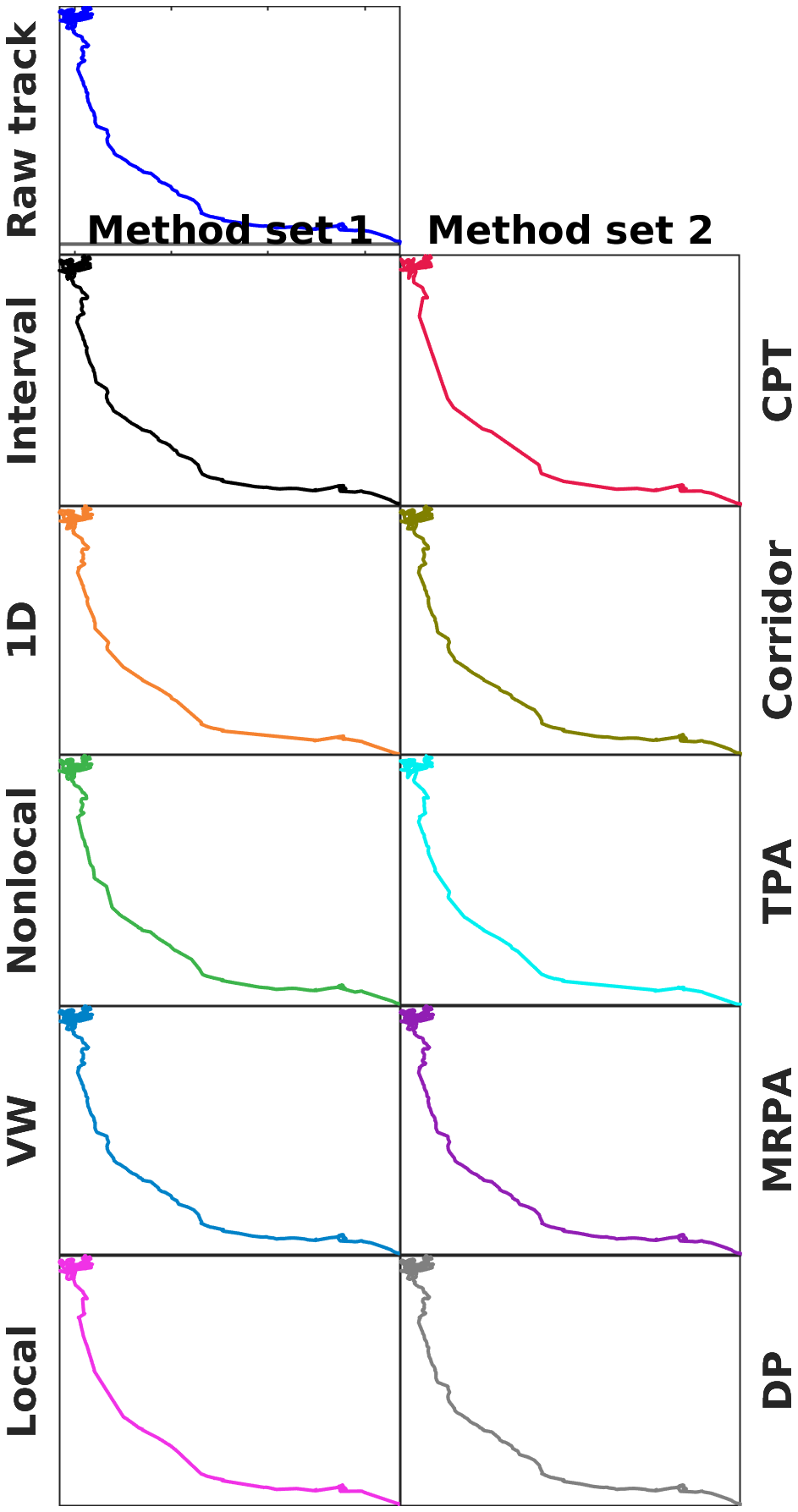

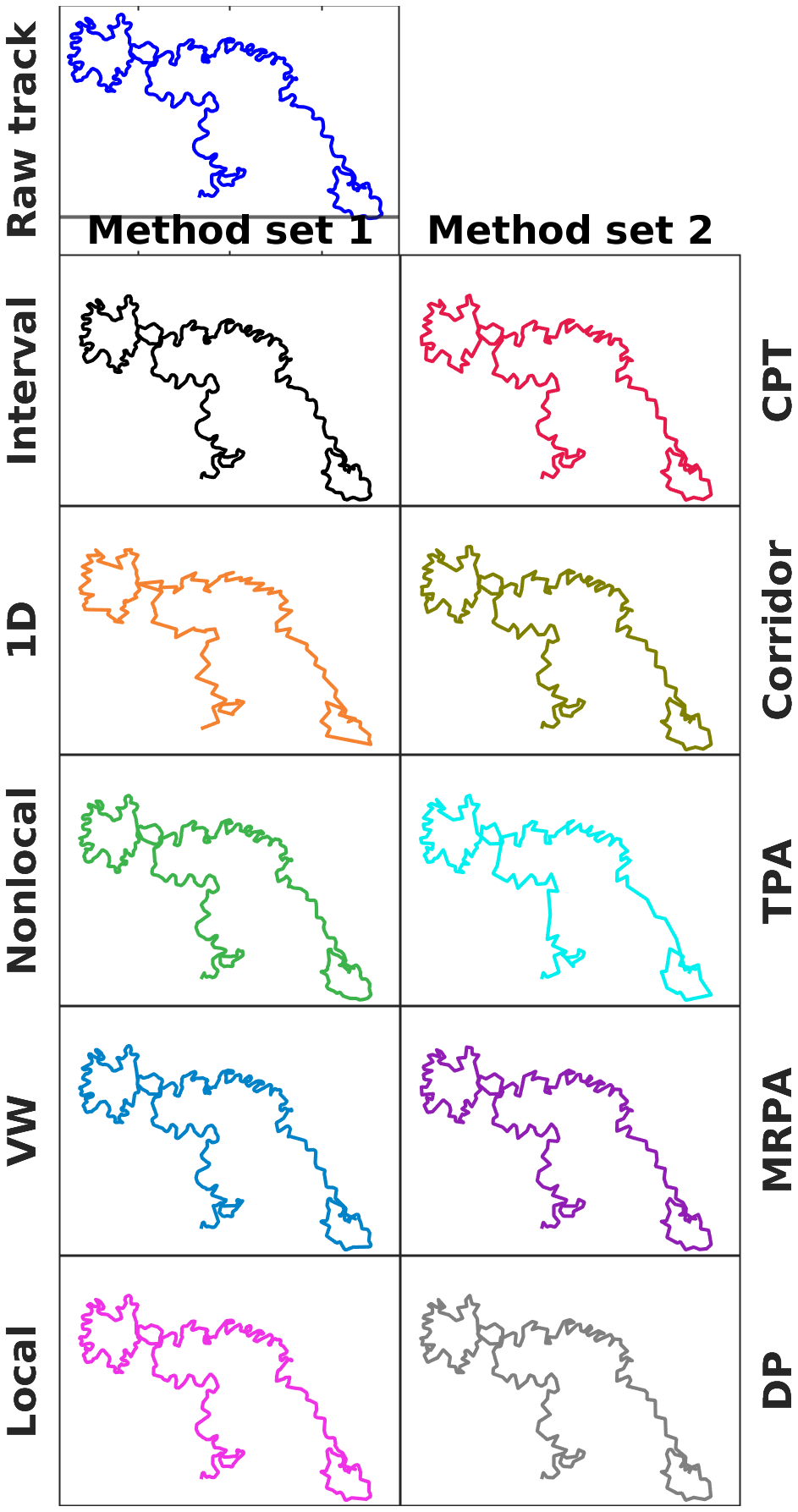

***Figure S2.*** ***Empirical resampled tracks****. Used track of a feral cat (*Felis catus*, left) and a rock ant (*Temnothorax rugatulus, *right) and tracks resulting from all tested resampling methods. Thresholds were chosen to select outcomes with the most similar mean step lengths.*

**Table** **S3**. All included resampling methods and their respective thresholds. Full names and descriptions of the methods can be taken from Table 1, and the selection process is explained in ‘Resampling’. Note that TPA and CPT use an additional threshold; we fixed the length parameter of TPA to the theoretically near-optimal mean step length and the *α* parameter of CPT to 0.06.

| **Method** | **Threshold** | | | | | | | | | |
| --- | --- | --- | --- | --- | --- | --- | --- | --- | --- | --- |
| **No.** | **1** | **2** | **3** | **4** | **5** | **6** | **7** | **8** | **9** | **10** |
| Interval | 2 | 4 | 7 | 13 | 18 | 24 | 31 | 39 | 48 | 58 |
| 1D | 0.001 | 0.005 | 0.01 | 0.1 | 1 | 5 | 10 | 50 | 100 | 500 |
| Nonlocal | 2 | 20 | 25 | 30 | 35 | 40 | 45 | 55 | 70 | 90 |
| VW | 0.2 | 0.4 | 0.6 | 1 | 1.5 | 2 | 3 | 4 | 8 | 16 |
| Local | 10 | 19 | 28 | 37 | 46 | 55 | 64 | 73 | 82 | 90 |
| CPT | 1 | 2 | 3 | 4 | 5 | 6 | 7 | 8 | 9 | 20 |
| Corridor | 0.1 | 0.6 | 1 | 1.4 | 1.8 | 2.2 | 2.6 | 3 | 3.4 | 5 |
| TPA | 0.001 | 5 | 10 | 15 | 20 | 25 | 30 | 35 | 40 | 100 |
| DP | 0.2 | 0.4 | 0.6 | 1 | 1.5 | 2 | 3 | 4 | 8 | 16 |
| MRPA | 0.25 | 5 | 10 | 50 | 100 | 500 | 1000 | 5000 | 10000 | 100000 |

**Table S4**. Methods which were not included in the comparisons in this paper because they include a speed parameter.

| **Method Name** | **Description** | **Citation** | **Used also in** |
| --- | --- | --- | --- |
| ‘vDP’ | 1. Track is projected to the space of: v·sin(θ), v·θ), with v = speed, θ = heading angle. 2. This projected trajectory is simplified with D-P (just above) and segments <3 points are regrouped | Thiebault et al. 2023 [1] | [1]: ARS, SSM, birds  [2]Tremblay [2013](https://journals.plos.org/plosone/article?id=10.1371/journal.pone.0088424): ST, FD, birds  [3]: env, birds  [4]: ST, α, env, birds  [5]: ST, SSM, env, birds  [6]: SSM, env, birds |
| STTrace | STTrace expands the Local method by adding a speed threshold, such that a point is kept if either the angle or the speed threshold is breached. | Potamias et al. [2006](https://ieeexplore.ieee.org/abstract/document/1644324?casa_token=GlhIwT-ZtE8AAAAA:iBLtJCB1JdJUBt8Rie8f5Sq5xcaVKg-p47b-TcbR1RQh_MUfjL8gJ2PH5iZJOEhlTf2FGGVR_Q) [7] | 240+ citations, Human movement |
| Viswalingam-Whyatt time-sensitive (VW-TS) | The smallest triangle formed between all sets of three consecutive points plotted in the time-space cube is iteratively dropped until its area is larger than the threshold | Hunnik [2017](https://studenttheses.uu.nl/handle/20.500.12932/25654) [8] | *Only citations in reviews and comparisons found.* |
| TraClus | Minimum Description Length cost of angle & distance (speed) | Lee et al. [2007](https://dl.acm.org/doi/abs/10.1145/1247480.1247546?casa_token=CSUW-YQNibYAAAAA:ISxA3uapPTwmCP-0WYAaEZUzgzyHbS4-GqnzomLbPQJOkHFVhgdZMSwB1fnbWmAJCSZOENsY0wB2) [9] | *Only citations in reviews and comparisons found.* |

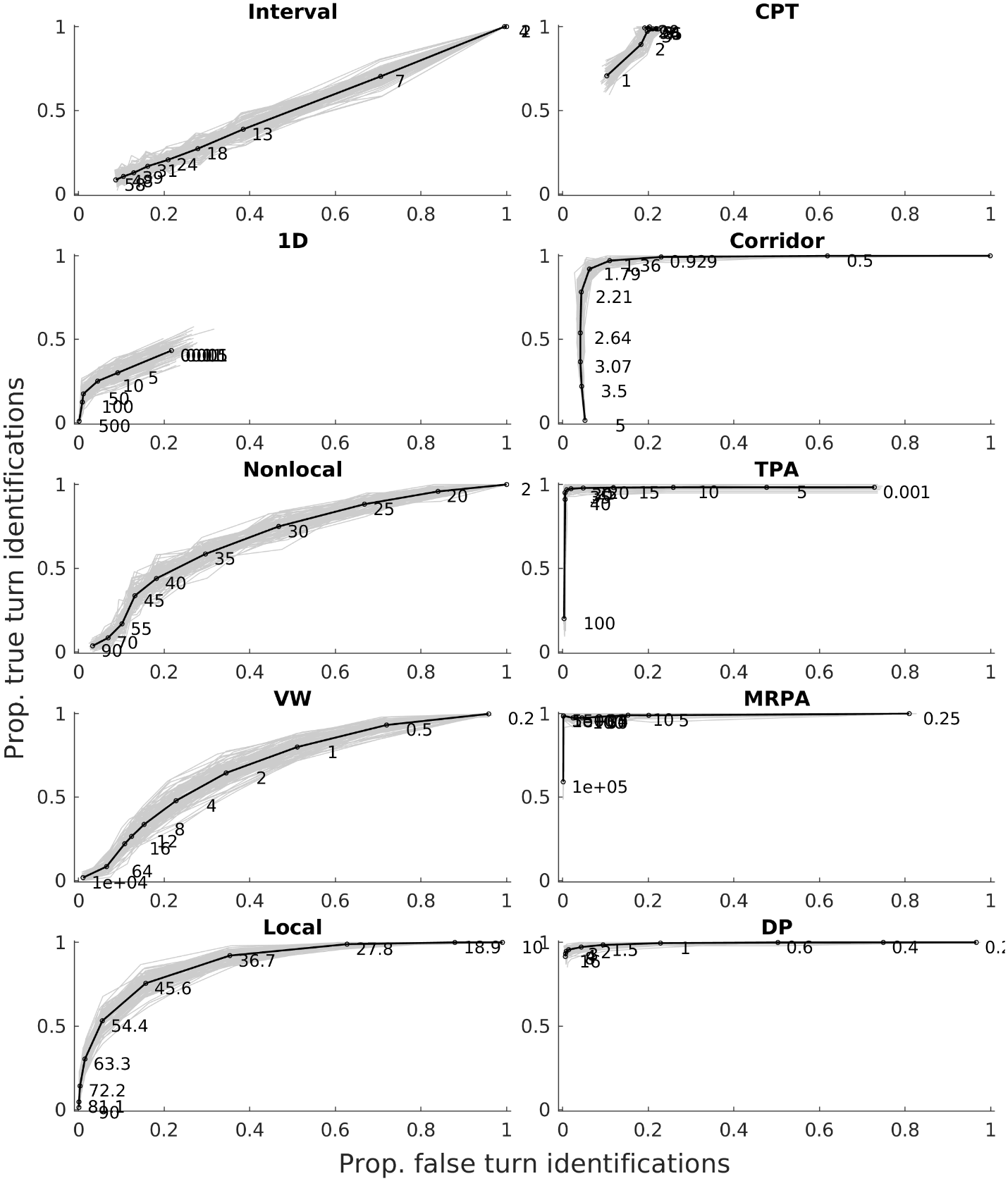

***Figure S5. All ROC curves for the default scenario.*** *Black lines are means and grey lines are resampled tracks of 100 replicate ground truth tracks per scenario.*

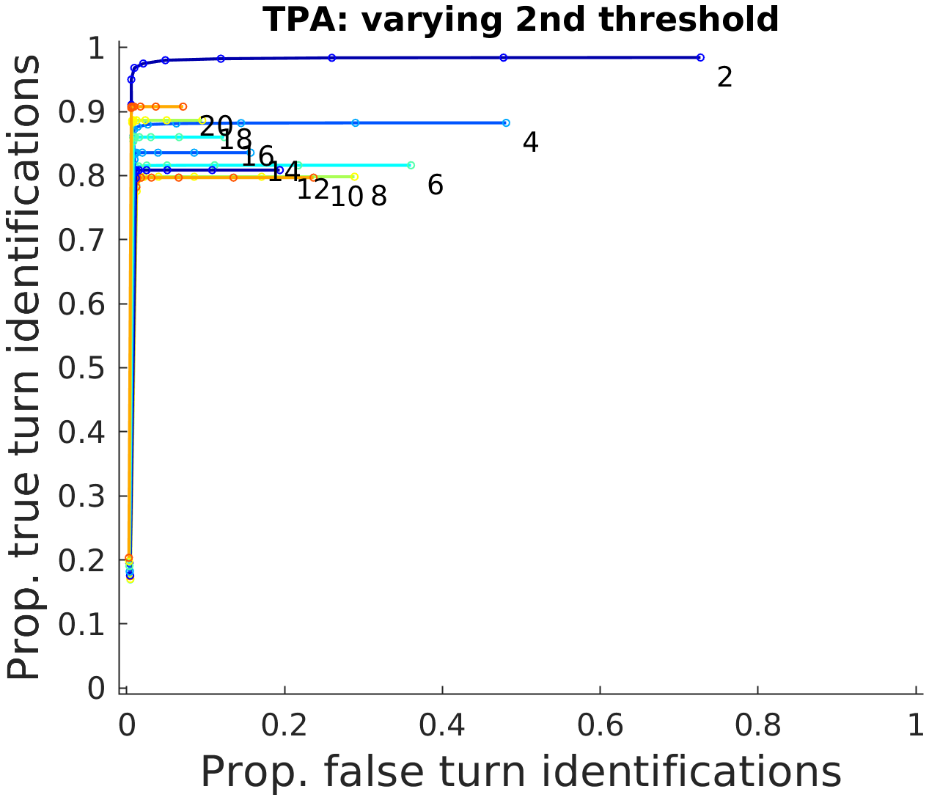

***Figure S6. ROC curve for TPA with alternative window size thresholds.*** *Different colored lines are ROC curves for different window sizes (i.e., the second threshold), indicated by the annotations. The first, angle-based thresholds, are the same as in the main analysis (see Table S3).*

### Results

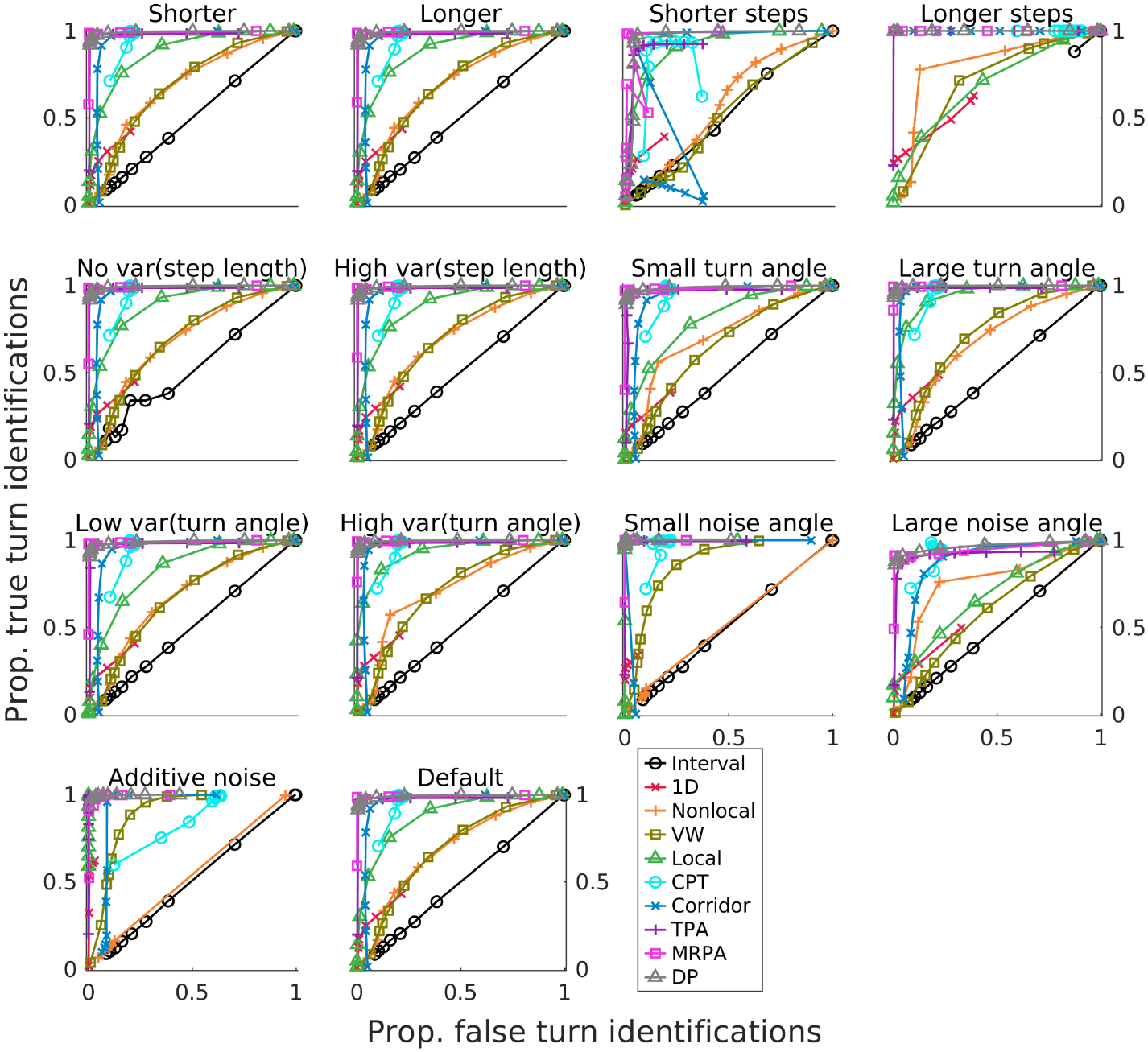
***Figure S7 ROC curves for all scenarios*** *(only means shown).*

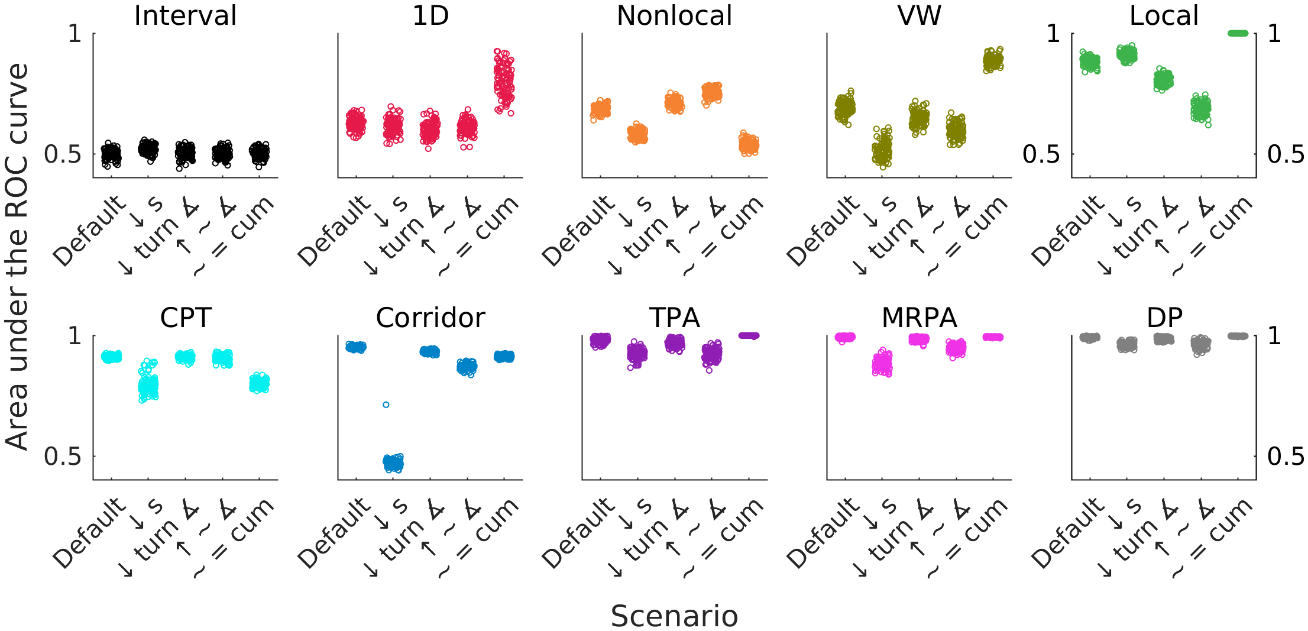

***Figure S8 Areas under the ROC curves*** *for the 5 main scenarios, grouped by method (instead of scenario, as in main text Figure 3).*

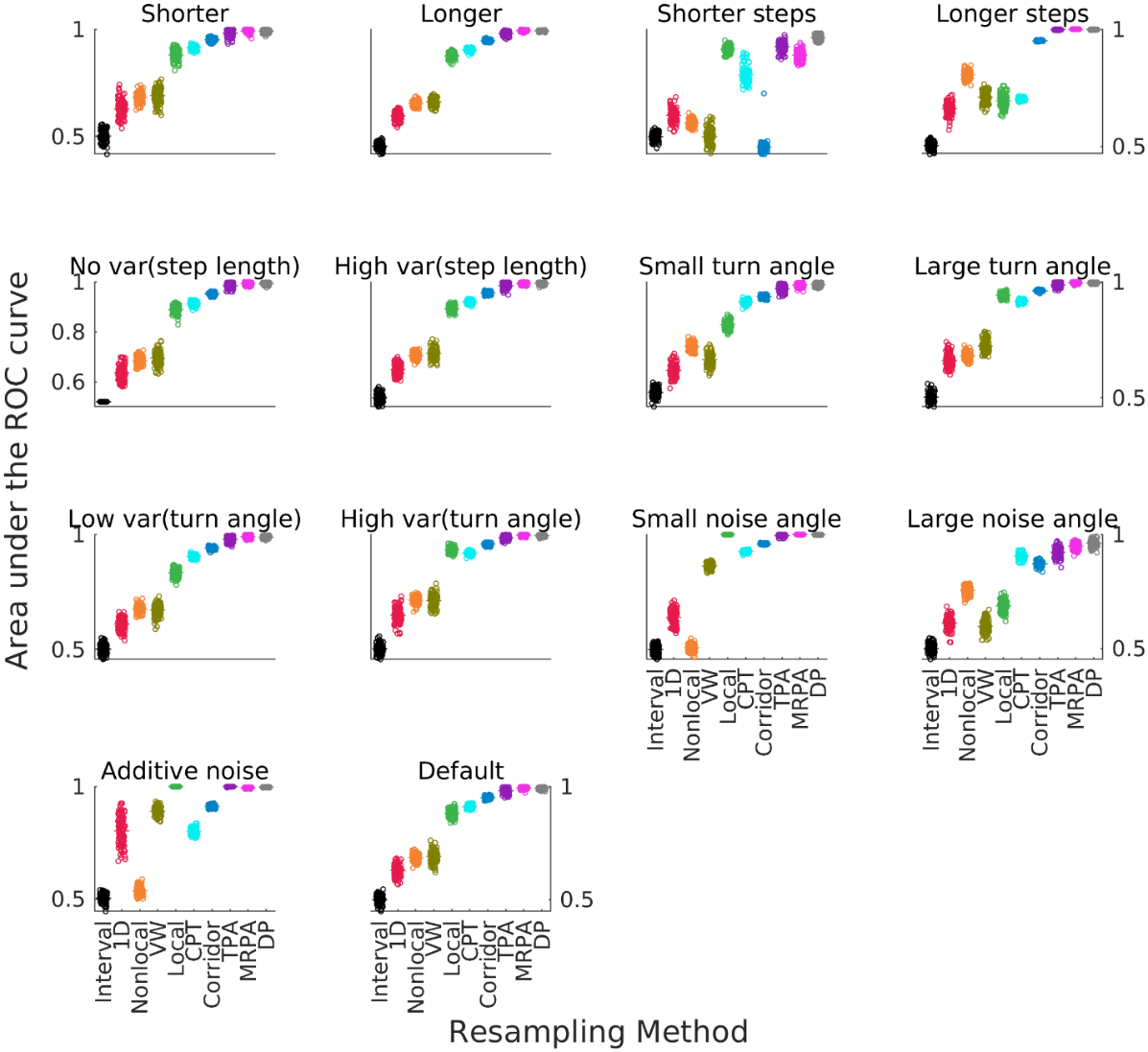

***Figure S9. All areas under the ROC curves****.* *Black horizontal lines are means and grey dots are resampled tracks of 100 replicate ground truth tracks per scenario. Larger numbers mean higher accuracy.*

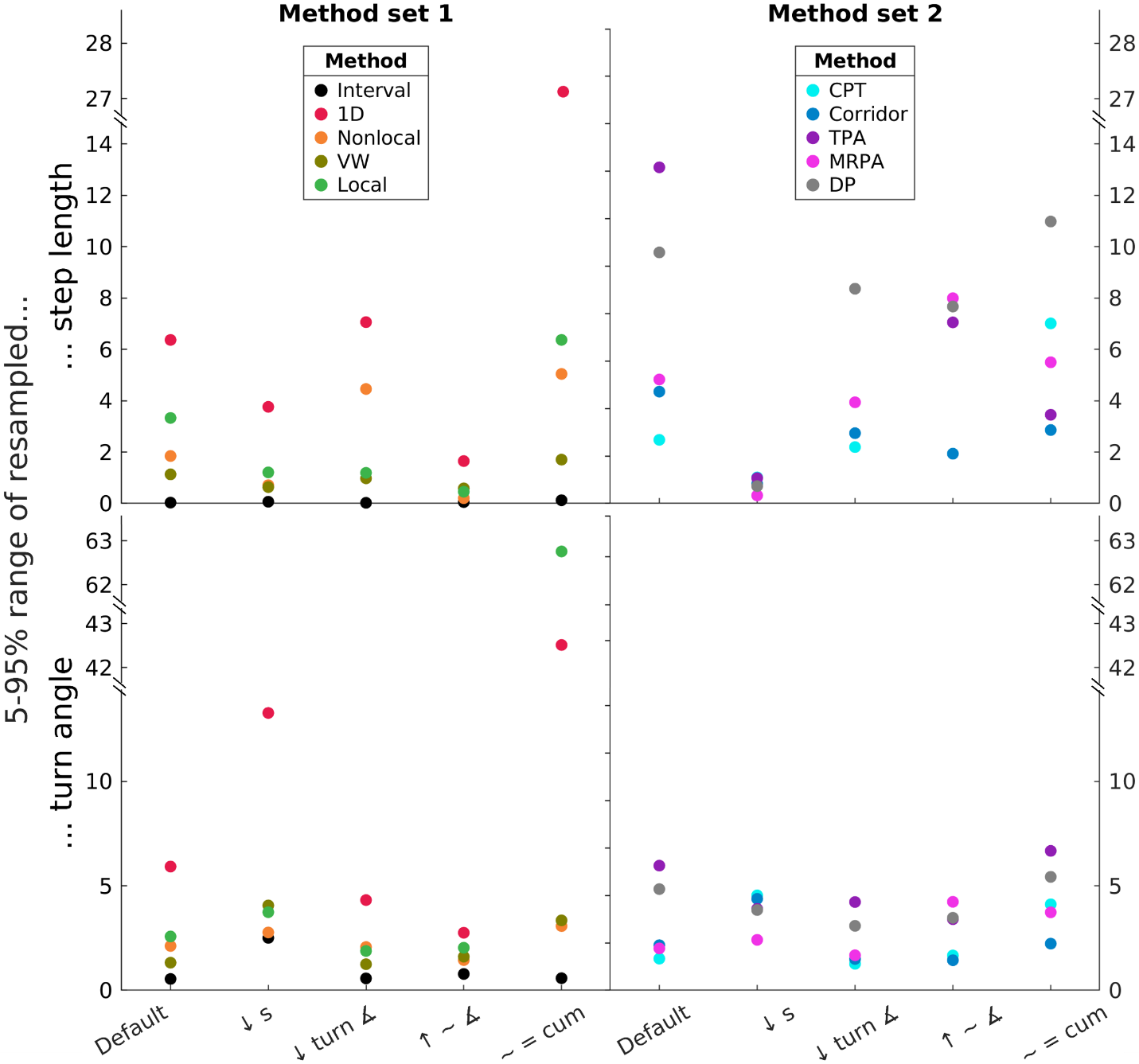

***Figure S10 Resampled track mean step length and mean absolute turn angle variability.*** *The x-axis shows the parameter sets shown in the previous figures: Default, Short steps, Small turn angle, Large noise angle, Cumulative noise. Different colors and symbols correspond to the different resampling methods. Left and right panels each show one half of the resampling methods for clarity. The y-axes display the range of the 5^th^ to 95^th^ percentile for a given set of 100 ground truth tracks of each scenario. For example, if 90 out of 100 resampled tracks had mean turn angles greater than 45° but smaller than 55°, this would result in a plotted range value of 10 (55-45). Perfect accuracy would result in a value of 0, since the calculated turn angle would always be the same for the same ground truth parameter. Top panels: step length, bottom panels: turn angle. All data are from resampled tracks resulting from the thresholds yielding the best turn identification accuracy as defined by the point which is closest to the (0,1) corner of the ROC curve, for the respective method.*
